# Genotypic Variation in Below-to Aboveground Systemic Induction of Glucosinolates Mediates Plant Fitness Consequences under Herbivore Attack

**DOI:** 10.1101/810432

**Authors:** Moe Bakhtiari, Sergio Rasmann

**Author notes:** Corresponding author: Moe Bakhtiari. Authors’ information: Sergio Rasmann.

## Abstract

Plants defend themselves against herbivore attack by constitutively producing toxic secondary metabolites, as well as by inducing them during herbivore feeding. Induction of secondary metabolites can cross plant tissue boundaries, such as from root to shoot. However, whether the potential for plants to systemically induce secondary metabolites from roots to shoots shows genetic variability, and thus, potentially, is under selection conferring fitness benefits to the plants is an open question. To address this question, we induced 26 maternal plant families of the wild species *Cardamine hirsuta* belowground (BG) using the wound-mimicking phytohormone jasmonic acid (JA). We measured resistance against a generalist (*Spodoptera littoralis*) and a specialist (*Pieris brassicae*) herbivore species, as well as the production of glucosinolates (GSLs) in plants. We showed that BG induction increased AG resistance against the generalist but not against the specialist, and found substantial plant family-level variation for resistance and GSL induction. We further found that the systemic induction of several GSLs tempered the negative effects of herbivory on total seed set production. Using a widespread natural system, we thus confirm that BG to AG induction has a strong genetic component, and it can be under positive selection by increasing plant fitness. We suggest that natural variation in systemic induction is in part dictated by allocation trade-offs between constitutive and inducible GSL production, as well as natural variation in AG and BG herbivore attack in nature.

## INTRODUCTION

The selective pressure of insect herbivores on plants has led to the evolution of a wide variety of secondary metabolites that can intoxicate or inhibit digestion capacities of the herbivores during feeding (Futuyma and Agrawal 2009; Schoonhoven et al. 2005). While secondary metabolites can be constitutively stored in plant tissues prior to herbivore attack, herbivore feeding on one organ of a plant can induce the *de novo* production, or increase accumulation of the toxins locally, on the same organ, or systemically, on other organs of a plant (Kessler and Baldwin 2002). Within-plant induction of toxic chemicals often reduces the performance of current or subsequent herbivores (Karban and Baldwin 1997; Poelman et al. 2008), and therefore, locally- or systemically-induced chemical defenses should be linked to plant fitness (Agrawal 1998; Agrawal 2000). Moreover, the induction of defenses can cross widely-separated plant organs, such as travelling from roots to shoots (Bezemer et al. 2003). Indeed, a growing body of literature is showing that a range of belowground (BG) organisms can induce defense responses in aboveground (AG) tissues and *vice versa* (Bezemer et al. 2003; Erb et al. 2009; Rasmann and Agrawal 2008; Soler et al. 2005; Staley et al. 2007; van Dam et al. 2005). Reviews on the topic suggest that the magnitude and direction of chemically-mediated AG-BG interactions in plants largely depend on plant genotypic variation as well as the attacking species’ identity (Kabouw et al. 2011; van Geem et al. 2013; Vandegehuchte et al. 2011). Therefore, potentially, selection is acting on plants’ natural genetic variation to optimize their ability to induce defenses systemically. While significant levels of genetic variation, as well as a heritable genetic basis for both constitutive and inducible defense traits expression has been shown in several systems (Agrawal et al. 2002; Havill and Raffa 1999; Humphrey et al. 2018; Stevens and Lindroth 2005; Underwood et al. 2000; Wagner and Mitchell-Olds 2018), to date, we have practically no information on whether BG-AG defense induction is under positive selection for plants harboring such trait variation in nature.. Measuring BG to AG root induction is also ecologically-relevant since in nature, plants can be potentially in contact with a wide range of root herbivores, that phenologically, can induce the plants before the leaf herbivores arrive (Erb et al. 2008; Huang et al. 2017; Rasmann and Agrawal 2008).

For a trait to be under selection, it needs to display a significant degree of genetically based variation in nature. Whereas most of such variation is generated by random mutation, and evolutionary and genetic mechanisms (Caliskan 2012), the maintenance of genetic variability can also be affected by energetic costs. Optimal defense theory suggests that inducible defenses have evolved as a cost-saving strategy, and the relative allocation of constitutive and inducible defenses in plant organs, individuals or populations depends on predictability of attack from herbivores, the susceptibility of plants to attacks, and the context dependency of the interaction (e.g. environmental variation) (Agrawal et al. 2002; McKey 1974; Zangerl and Bazzaz 1992; Zangerl and Rutledge 1996). In other words, the simultaneous expression of constitutive and induced defense is thought to be costly (Brody and Karban 1992; Rasmann and Agrawal 2009; Strauss et al. 2002; Zangerl and Bazzaz 1992) and should result in negative genetic correlations (trade-offs) between individual traits and between defense deployment strategies (Agrawal et al. 2010). Therefore, high constitutive expression of a defense trait is predicted to be associated with lower induction abilities. While trade-offs between constitutive and induced defenses on the same organs have been shown in several systems (Heil et al. 2004; Rasmann et al. 2015; Rasmann et al. 2011; Thaler et al. 1999; Thaler and Karban 1997), we still lack evidence for whether AG inducibility of defenses after BG induction is trading-off with constitutive defenses.

In brassicaceaous plants, glucosinolates (GSLs), Sulphur- and nitrogen-containing plant secondary metabolites, are the main defensive compounds conferring plant resistance against insect herbivores (Howe and Jander 2008). The defensive function of GSLs breakdown products, when either expressed constitutively or when induced, against both specialist and generalist insect herbivores has been amply documented (Agrawal 1998; Agrawal 2000; Agrawal et al. 2002; Baldwin 1998; Karban and Baldwin 1997; van Dam and Oomen 2008; van Dam and Raaijmakers 2005; van Dam et al. 2005). Several individual GSLs show strong inducibility following herbivory and generally, the plant hormone jasmonic acid (JA) is a key player in the regulation of induced plant responses against chewing herbivores such as caterpillars (Farmer et al. 2003; Howe and Jander 2008). Emerging patterns from studies on *Brassica* spp. indicate that BG insect herbivory, or JA application to roots increase total GSLs levels in shoots (Griffiths et al. 1994; Pierre et al. 2012; Soler et al. 2005; van Dam et al. 2004). For instance, previous work with *Cardamine hirsuta* demonstrated that the overall abundance and identity of GSLs in the leaves is affected by JA induction in the roots (Bakhtiari et al. 2018). Therefore, if genetic variation for root-to-shoot induction exists in nature, it should correlate to plant fitness, particularly, when plants are under herbivore attack.

We here sought for natural genetic variability in BG-to-AG systemic induction in nature and specifically asked the following questions: 1) Does the exogenous application of JA in roots increase resistance against specialist and generalist insect leaf-chewing herbivores? 2) Is there genetically-based variation in resistance against insects and BG-to-AG induction of GSLs? 3) Is there a trade-off between the constitutive and inducible production of shoot GSLs following root induction? and 4) What is the impact of systemic induction of different GSLs on plant fitness? We answered these questions by inducing the roots of 26 maternal half-sib families of *Cardamine hirsuta* (Brassicaceae), measured GSL production in the leaves, and measured the growth of a specialist herbivore, the large cabbage butterfly *Pieris brassicae*, and a generalist noctuid butterfly, *Spodoptera littorali*s, to assess the potential impact of GSLs on adapted and non-adapted herbivores, respectively Our work builds toward a better understating of the ecological and evolutionary drivers of plant chemical defense variation in nature.

## METHODS AND MATERIALS

### Plants and Insects

The hairy bittercress, *Cardamine hirsuta* (Brassicaceae) is a common weed growing in a variety of habitats in Europe but mainly at low elevations (Pellissier et al. 2016). Seeds from 26 half-sib families were collected from three different natural populations separated by at least 10 Km (pop A = 9 pop B = 10, and pop C = 7 families) at the foothills of the Swiss Jura mountains. After an overwintering period of four months at 4 °C, seeds were germinated in Petri dishes lined with humid filter paper, and one week after germination, 15 seedlings per family (total of 390 plants) were transplanted independently into plastic potting pots (13 cm width × 10 cm height) filled with 500 ml of sieved soil (1 cm mesh size) mixed with sand in a 3:1 ratio. The soil/sand mixture was sterilized by autoclave at 120 °C for four hrs. Plants were immediately transferred to climate-controlled chamber and kept at 16h/22°C - 8h/16°C day-night, and 50% relative humidity conditions. Plants were fertilized (universal liquid fertilizer containing N: P: K ratio of 7:3:6% per liter) twice a week until the beginning of experiment. Although our common garden experiment was specifically designed to measure genetic variation across different maternal lines, we would like to acknowledge that the potentially observed genetic differences among families cannot be fully isolated from maternal environmental effects, but because *C. hirsuta* practically completely relies on autogamous selfing for reproduction (Hay et al. 2014), such maternal effects should be minimized in this system. We used the large cabbage butterfly *Pieris brassicae* (Lepidoptera: Pieridae) and the African cotton leaf worm *Spodoptera littoralis* (Lepidoptera: Noctuidae) as specialist and generalist herbivore insects, respectively. *P. brassicae* is a specialist herbivore that feeds exclusively on plants producing GSLs, especially on species of the Brassicaceae (Chew 1988). The caterpillars used in this experiment were obtained from a culture maintained on *Brassica rapae* ssp. *chinensis* (L) plants. *S. littoralis* is a generalist herbivore known to feed on species belonging to more than forty families of plants (Brown and Dewhurst 1975). However, it does not occur in Switzerland, therefore, it functioned as a generalist, non-adapted, herbivore in our study. Eggs were obtained from Syngenta, Stein AG, Switzerland, and newly hatched *S. littoralis* larvae to be used in the bioassays were reared on corn-based artificial diet until the beginning of the experiment.

### Experimental Design and Insect Bioassay

After three weeks of growth, we randomly assigned the plants to three treatment groups. Six plants per family were randomly assigned to the JA treatment, another six group plants to the no-induction treatment, and the rest (three plants per family) to the no-herbivory control treatment. Each plant in the JA treatment were inoculated with 20 ml of JA solution in the roots by spiking the solution into the soil, 0.5 cm below the surface. The JA solution consisted of 2.4 µmoles (500 µg) of JA (± - jasmonic acid, Sigma, St Louis, IL, USA) per plant in 10 ml demineralized water and 0.5% EtOH (van Dam and Oomen 2008; van Dam et al. 2004). The no-induction group of plants received 20 ml of 0.5% EtOH in acid water (pH 3.7 with HCl). We chose to induce roots with JA instead of using a root herbivore (e.g. cabbage root maggots), in order to standardize the induction event across all plant families. Moreover, by applying JA, we intentionally avoided the effect of tissue removal per se on plant fitness. In other words, we were able to measure the fitness impact of defense induction independently from herbivore damage.

To measure the effect of BG induction on leaf chemistry, four days after JA root application, we collected two fully-expanded new leaves per plant in the JA and the no-induction treatments, and immediately froze and stored them at −80°C for further chemical analyses. Since the leaves from both treatments were collected prior to AG herbivory, the plant materials collected from no-induction treatment served for measuring constitutive secondary metabolites expression. Immediately after leaf removal, we infested half of the plants in the herbivory treatments (three plants per family per induction treatment = six plants/family) with two 7-days old *S. littoralis* larvae, and the other half, with one 6-days old *P. brassicae* larvae. We next covered all plants with gauze bags to prevent escape or cross-movement of insects between plants. After one week of herbivory, bags were removed, the insects were retrieved from individual plants, and their combined weight per plant was measured and recorded to obtain the average insect weight per plant. We used the formula ln(*final weight*− *initial weight*) to determine the insects’ weight gain and plant resistance (i.e. lower growth rate indicate that plants are more resistant). After the herbivore bioassay, we allowed the plants to complete their life cycle and produce seeds. To estimate the total seed production on each plant, we first randomly selected one seedpod per plant from 50 plants, measured each pod’s length, and counted the number of seeds per pod. Using these data, we fitted a linear regression of the seed number as the function of seedpod length in order to obtain the seed set of each plants based on the seedpod length (equation: 14.92 × *total pods length* + 1.65). At the end of the experiment, when all seedpods were mature, AG plant parts were separated from roots, oven-dried at 40°C for 48h and weighted to determine their dry biomass, which served as covariate in the statistical analyses (see below).

### Glucosinolate Analyses

We assessed the concentration of individual GSLs in leaf tissues in no-induction and root-JA-induction plants prior to the AG herbivore application. This allowed measuring the chemical content of the leaves to which the herbivores were immediately exposed across different treatments, and to measure the direct effect of the root induction treatment on plant chemistry without the confounding effect of additional herbivore feeding. To this end, we ground the fresh leaves to powder using mortars and pestles in liquid nitrogen. A 100-mg aliquot of fresh leaf powder was then added with 1.0 ml Methanol: H2O: formic acid (80:19.5:0.5, v/v) and 5 glass beads in Eppendorf tubes, shaken in a tissuelyser (Mixer Mill MM 400, Retsch GmbH, Haan, Germany) for 4 min at 30 Hertz, and centrifuged them at 12800 g for 3 min. The supernatant was then transferred to HPLC vials for liquid chromatography analysis. Glucosinolate identification and quantification was performed using an Acquity UPLC from Waters (Milford, MA, USA) interfaced to a Synapt G2 QTOF from Waters with electrospray ionization, using the separation and identification method as described in Glauser et al. (2012).

### Statistical Analyses

All statistical analyses were carried out with R software (R Development Core Team 2017).

#### 1) Does the exogenous application of JA in roots increase resistance against specialist and generalist insect leaf-chewing herbivores?

To answer this question we performed two ANOVAs on the larval weight gain, for generalist and specialist respectively, with the JA treatment (two levels) as fixed factor.

#### 2) Is there genetically-based variation in insect resistance and BG-to-AG GSLs induction?

First, we assessed the effect of JA treatment (two levels) and maternal families (26 families) on the abundance and composition of all GSLs simultaneously using a permutational multivariate analysis of variance (PERMANOVA) with the *adonis* function in the package *vegan* (Oksanen et al. 2017). We included plant biomass as covariate to control for potential direct effect of plant size (Züst et al. 2015) on GSL production, and populations as strata in the model. The results were visualized using a non-metric multidimensional scaling (NMDS) ordination. The Bray–Curtis metric was used to calculate a dissimilarity matrix of all compounds among samples for both the PERMANOVA and the NMDS. Second, to address the effect of root JA-addition and family variation on all i) individual GSLs production, ii) the total amount of GSLs, iii) the AG resistance against *P. brassicae* and *S. littoralis* (insect weight gain), iv) and the seed production, we ran linear mixed-effect models with JA treatment as fixed factor, plant families nested within populations as random factor, and plant biomass as covariate, using the function *lme* in the package *nlme* (Pinheiro et al. 2017). Because families were included as random factor in the initial model, we estimated their effect by running a second model without the nested family factor. Differences between the first and the second models (AIC scores) would inform on potential maternal family variation, which were assessed using log-likelihood ratios and Chi-Square tests (function *Chisquare* in R). In addition, to test for family-level genetic variation in inducibility (G x E) on all individual GSLs and total amount of GSLs per plant, we ran ANCOVAs with JA treatment, plant families nested within populations and their interactions as fixed factors, and plant biomass as covariate, using the function *lm* in R.

#### 3) Is there a trade-off between the constitutive and inducible production of shoot GSLs following root induction?

To test for trade-offs between the constitutive production and the inducibility of total GSLs among the 26 plant families, we employed a Monte Carlo simulation procedure proposed by Morris et al. (2006) using MATLAB (Version 7.5.0.342 –R2007b, MathWorks Inc., USA). This statistical approach accounts for several issues that have apparently confounded previous attempts to assess a trade-off between constitutive and induced defenses (Morris et al. 2006). Specifically, this approach uses the difference in mean GSL production between JA-treated and control plants for measuring induced production of GSLs, and uses a modified Monte Carlo procedure that takes into account sampling variation due to limited sample size, measurement error from environmental and genetic differences.

#### 4) What is the impact of systemically inducing different GSLs on plant fitness?

First, we tested for the effect of treatment (3 levels in this case: root JA induction, no-induction, no-herbivory control treatment) on lifetime seed production using mixed effect models with JA treatment as fixed factor and families nested in populations as random factor, including biomass as covariate (*lme* function), followed by pairwise comparisons using *Tukey HSD* post-hoc tests (*lsmeans* function in the package *lsmeans* (Lenth 2016). Second, to estimate the lifetime fitness effect of root JA induction across all different GSLs, we ran mixed effect ANCOVA models with seed production per plant as response variable, individual and total GSLs in interaction with JA induction treatment as continuous and categorical fixed factors in the model, respectively. Plant families nested within population were included as random factor using the function *lme* in the package *nlme.* The aim here was to detect a significant interaction between JA induction and GSLs on seed set (as a proxy for plant fitness). If this were the case, it would indicate that the effect of JA treatment on a particular GSL compound would affect plant fitness, positively or negatively.

## RESULTS

### Effect of JA Treatment on Insect Resistance

We found that *S. littoralis* larvae on JA-treated plants grew 47% less compared to control plants (Fig. 1, Table 1), and maternal families responded differently in resistance against this generalist herbivore (Fig. 2a, Table 1). In contrast, *P. brassicae* larval weight gain did not differ between treatments and there was no family effect on larval weight gain (Table 1).

**Table 1.** Mixed effect model table for testing the effect of JA induction treatment in the roots of *Cardamine hirsuta* plants, maternal families, and their biomass on individual and total glucosinolates (GSL*), as well as seed production and *Spodoptera littoralis* larval growth. *C. hirsuta* plant families nested within populations was the random factor. Family effect was calculated from the log-likelihood difference (LLR) between the full model and the model missing the random effect. 16 GSL out of 28 showed a significant Family effect. GSL 9, 12, 14, 17, 26 showed significant JA effect. GSL 11, 24, 25, 26 showed significant biomass effect

| GSL † | Factor | Value | Df | t-value/LLR | p-value |  |
| --- | --- | --- | --- | --- | --- | --- |
| GSL1 | JA | 0.11 | 91 | 1.154 | 0.252 |  |
|  | Plant biomass | 0 | 91 | -0.896 | 0.373 |  |
|  | Family |  | 4 | 1.735 | 0.187 |  |
| GSL2 | JA | -0.062 | 91 | -1.19 | 0.237 |  |
|  | Plant biomass | 0 | 91 | 0.055 | 0.956 |  |
|  | Family |  | 4 | 0 | 1 |  |
| GSL3 | JA | 0.063 | 91 | 0.88 | 0.381 |  |
|  | Plant biomass | 0 | 91 | 0.769 | 0.444 |  |
|  | Family |  | 4 | 12.671 | 0 | *** |
| GSL4 | JA | 0.047 | 91 | 0.433 | 0.666 |  |
|  | Plant biomass | 0 | 91 | 1.362 | 0.177 |  |
|  | Family |  | 4 | 51.395 | 0 | *** |
| GSL5 | JA | 0.036 | 91 | 0.394 | 0.695 |  |
|  | Plant biomass | 0 | 91 | 0.673 | 0.502 |  |
|  | Family |  | 4 | 6.591 | 0.01 | ** |
| GSL6 | JA | 0.025 | 91 | 0.15 | 0.881 |  |
|  | Plant biomass | 0 | 91 | -0.104 | 0.918 |  |
|  | Family |  | 4 | 0.099 | 0.753 |  |
| GSL7 | JA | 0 | 91 | -0.005 | 0.996 |  |
|  | Plant biomass | 0 | 91 | 0.118 | 0.907 |  |
|  | Family |  | 4 | 7.301 | 0.006 | ** |
| GSL8 | JA | 0.047 | 91 | 0.284 | 0.777 |  |
|  | Plant biomass | 0.001 | 91 | 1.491 | 0.139 |  |
|  | Family |  | 4 | 18.37 | 0 | *** |
| GSL9 | JA | -0.086 | 91 | -2.846 | 0.006 | ** |
|  | Plant biomass | 0 | 91 | -0.299 | 0.766 |  |
|  | Family |  | 4 | 27.569 | 0 | *** |
| GSL10 | JA | -0.02 | 91 | -0.297 | 0.768 |  |
|  | Plant biomass | 0 | 91 | 0.33 | 0.742 |  |
|  | Family |  | 4 | 3.254 | 0.071 | ° |
| GSL11 | JA | 0.094 | 91 | 1.178 | 0.242 |  |
|  | Plant biomass | 0 | 91 | 2.168 | 0.033 | ** |
|  | Family |  | 4 | 19.56 | 0 | *** |
| GSL12 | JA | -0.095 | 91 | -1.769 | 0.08 | ° |
|  | Plant biomass | 0 | 91 | 0.808 | 0.421 |  |
|  | Family |  | 4 | 3.021 | 0.082 | ° |
| GSL13 | JA | 0.036 | 91 | 0.605 | 0.547 |  |
|  | Plant biomass | 0 | 91 | 1.476 | 0.143 |  |
|  | Family |  | 4 | 21.435 | 0 | *** |
| GSL14 | JA | -0.112 | 91 | -2.165 | 0.033 | ** |
|  | Plant biomass | 0 | 91 | 0.046 | 0.963 |  |
|  | Family |  | 4 | 3.043 | 0.081 | ° |
| GSL15 | JA | -0.081 | 91 | -1.019 | 0.311 |  |
|  | Plant biomass | 0 | 91 | -0.517 | 0.606 |  |
|  | Family |  | 4 | 0.031 | 0.859 |  |
| GSL16 | JA | 0.062 | 91 | 0.884 | 0.379 |  |
|  | Plant biomass | 0 | 91 | 0.172 | 0.864 |  |
|  | Family |  | 4 | 1.783 | 0.181 |  |
| GSL17 | JA | -0.094 | 91 | -1.74 | 0.085 | ° |
|  | Plant biomass | 0 | 91 | 0.754 | 0.453 |  |
|  | Family |  | 4 | 2.751 | 0.097 |  |
| GSL18 | JA | 0.021 | 91 | 0.126 | 0.9 |  |
|  | Plant biomass | 0 | 91 | 0.44 | 0.661 |  |
|  | Family |  | 4 | 0 | 1 |  |
| GSL19 | JA | -0.092 | 91 | -1.71 | 0.091 |  |
|  | Plant biomass | 0 | 91 | 0.778 | 0.439 |  |
|  | Family |  | 4 | 2.579 | 0.108 |  |
| GSL20 | JA | -0.057 | 91 | -1.535 | 0.128 |  |
|  | Plant biomass | 0 | 91 | -0.393 | 0.696 |  |
|  | Family |  | 4 | 0.857 | 0.005 | ** |
| GSL21 | JA | -0.091 | 91 | -1.768 | 0.081 |  |
|  | Plant biomass | 0 | 91 | -0.052 | 0.959 |  |
|  | Family |  | 4 | 2.735 | 0.098 | ° |
| GSL22 | JA | 0.113 | 91 | 0.804 | 0.424 |  |
|  | Plant biomass | 0 | 91 | -0.228 | 0.821 |  |
|  | Family |  | 4 | 14.837 | 0 | *** |
| GSL23 | JA | -0.091 | 91 | -1.768 | 0.081 | ° |
|  | Plant biomass | 0 | 91 | -0.052 | 0.959 |  |
|  | Family |  | 4 | 2.735 | 0.098 | ° |
| GSL24 | JA | 0.069 | 91 | 0.634 | 0.528 |  |
|  | Plant biomass | -0.001 | 91 | -4.379 | 0 | *** |
|  | Family |  | 4 | 0.812 | 0.367 |  |
| GSL25 | JA | -0.009 | 91 | -0.081 | 0.935 |  |
|  | Plant biomass | -0.002 | 91 | -5.913 | 0 | *** |
|  | Family |  | 4 | 3.536 | 0.06 | ° |
| GSL26 | JA | 0.04 | 91 | 1.888 | 0.062 | ° |
|  | Plant biomass | 0 | 91 | -3.367 | 0.001 | ** |
|  | Family |  | 4 | 9.61 | 0.002 | *** |
| GSL27 | JA | -0.089 | 91 | -1.649 | 0.103 |  |
|  | Plant biomass | 0 | 91 | 0.199 | 0.843 |  |
|  | Family |  | 4 | 1.311 | 0.252 |  |
| GSL28 | JA | -0.109 | 91 | -2.082 | 0.04 |  |
|  | Plant biomass | 0 | 91 | -0.027 | 0.978 |  |
|  | Family |  | 4 | 1.966 | 0.161 |  |
| GSL Total | JA | 0.029 | 91 | 0.194 | 0.847 |  |
|  | Plant biomass | 0 | 91 | 0.489 | 0.626 |  |
|  | Family |  | 4 | 0 | 1 |  |
| Seed production | JA | -80.991 | 116 | -1.697 | 0.092 | ° |
|  | Plant biomass | 0.171 | 116 | 1.206 | 0.23 |  |
|  | Family |  | 4 | 19.551 | 0 | *** |
| *S. littoralis* | JA | -0.835 | 92 | -4.736505 | 0 | *** |
|  | Plant biomass | 0 | 92 | -0.087 | 0.93 |  |
|  | Family |  | 4 | 13.989 | 0 | *** |
| *P. brassicae* | JA |  | 92 |  | 0.2 |  |
|  | Plant biomass |  | 92 |  | 0.4 |  |
|  | Family |  | 4 |  | 0.32 |  |

Signif. codes: \*\*\* < 0.001, \*\* < 0.01, \* < 0.05, ° < 0.1
† GSL1 = Glucoraphanin; GSL2 = Hydroxypropyl gsl; GSL3 = Progoitrin; GSL4 = Glucoalyssin; GSL5 = Glucoputranjivin; GSL6 = Gluconapin; GSL7 = Butyl gsl; GSL8 = Glucobrassicinapin; GSL9 = Glucohirsutin; GSL10 = Glucoerucin; GSL11 = Glucoberteroin; GSL12 = 8-Methylthiooctyl gsl ; GSL13 = Gluconapoleiferin; GSL14 = Hydroxymethylbutyl gsl ; GSL15 = 2-Methylbutyl gsl; GSL16 = Sinalbin; GSL17 = Veratryl gsl; GSL18 = Glucotropeolin; GSL19 = Trimethoxy gsl; GSL20 = 5-Benzoyloxypentyl ; GSL21 = 2-Hydroxy-2-phenylethyl gsl; GSL22 = Gluconasturtiin; GSL23 = Hydroxybenzyl-methylether gsl; GSL24 = Glucobrassicin; GSL25 = Methoxyglucobrassicin; GSL26 = Neoglucobrassicin; GSL27 = Unknown.C16H23NO10S2; GSL28 = Unknown.C19H28N3O12S3.

**Fig. 1.**
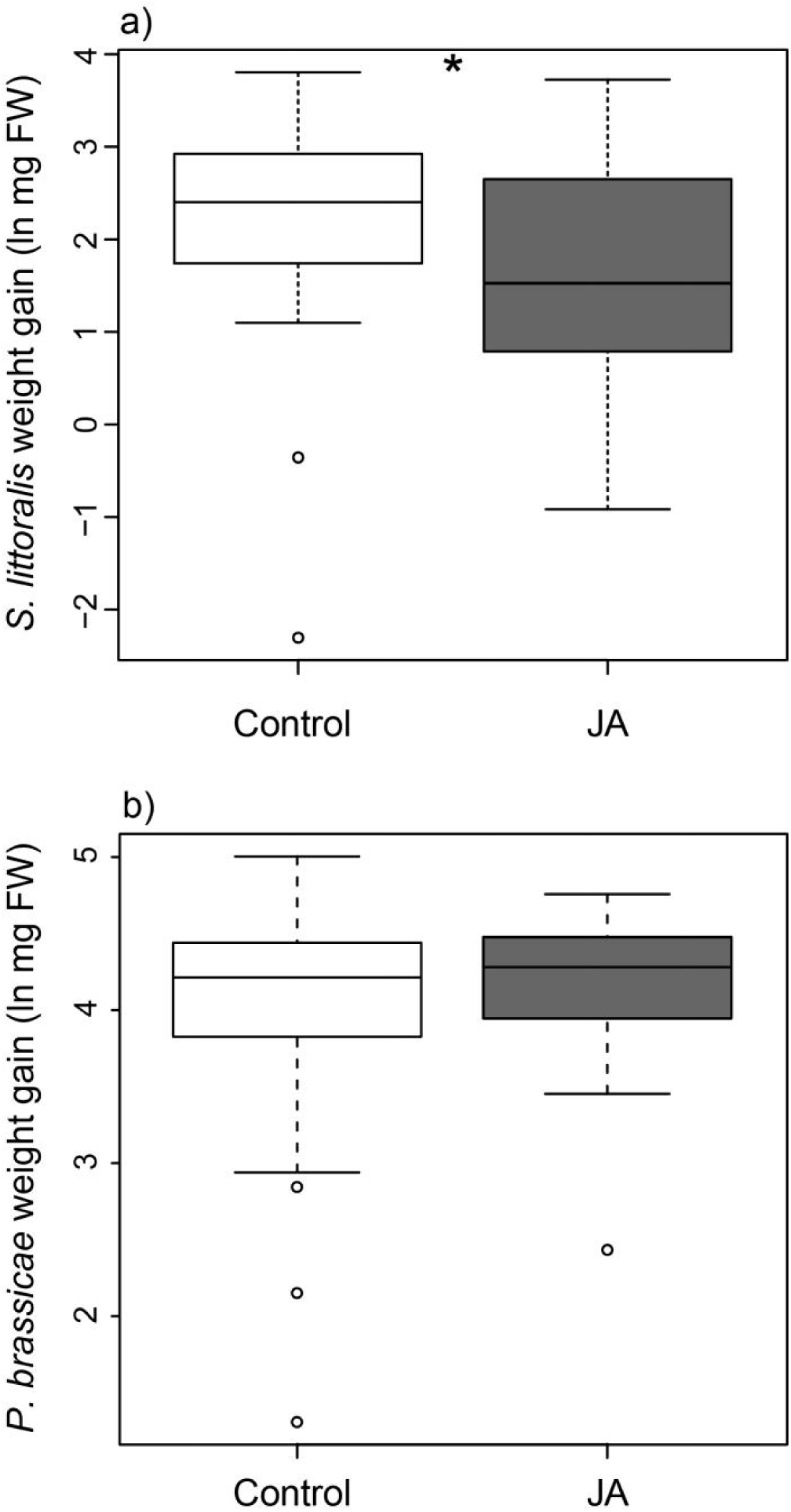
Root to shoot induction of resistance against generalist and specialist herbivores. Boxplots show average weight gain of (a) *Spodoptera littoralis*, and (b) *Pieris brassicae* caterpillars feeding on plants that received jasmonic acid (JA) in the roots 4 days prior herbivory (JA, grey boxes), or received no JA in the roots (Control, open boxes)). Weight gain was calculated as the natural logarithm of the difference between final and initial fresh weight. Asterisks show significant differences across the two treatments (p < 0.05).

**Fig. 2.**
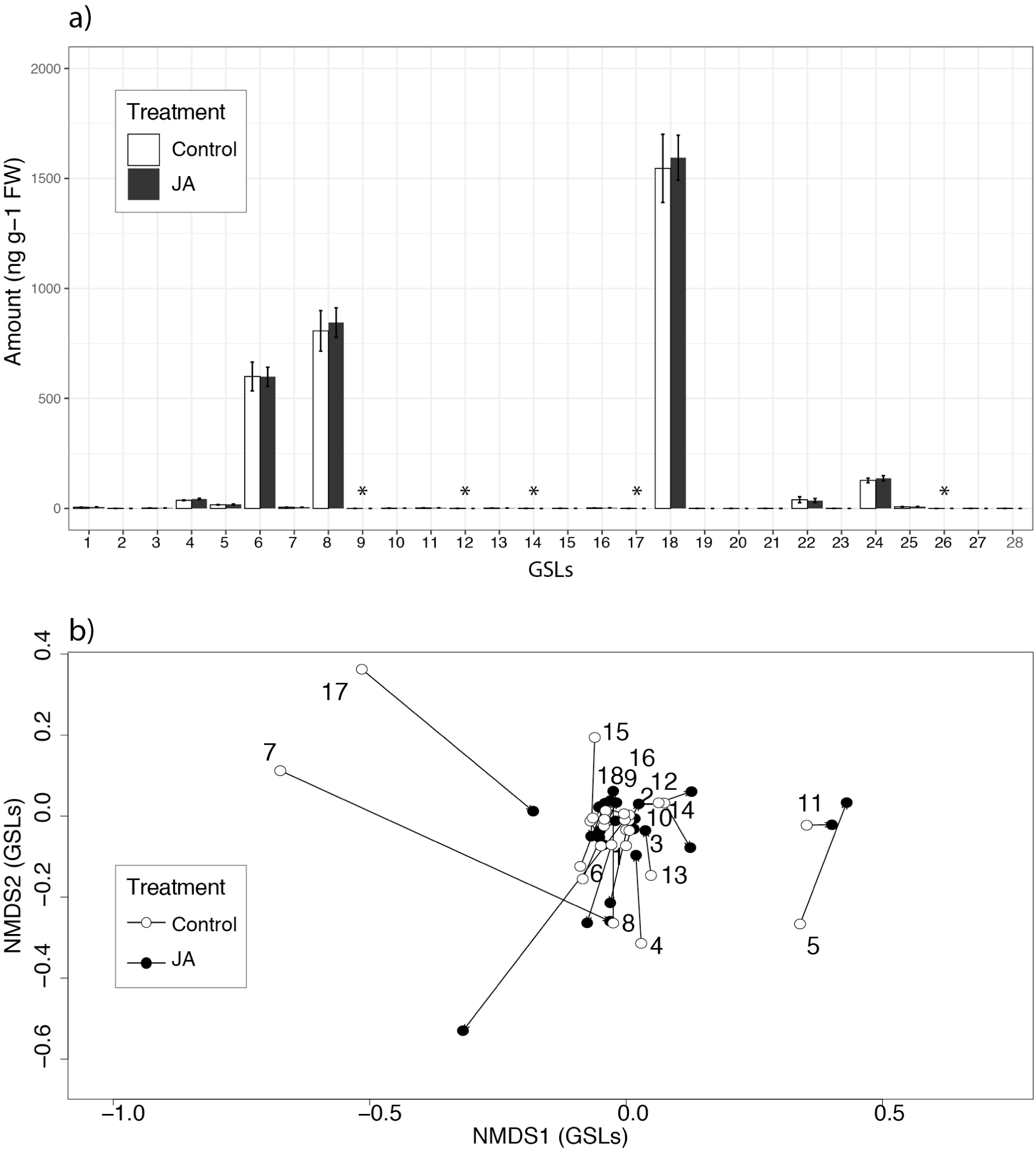
Genetic variation in resistance and fitness related traits. Shown are reaction norm plots for the effects of JA treatment in roots on a) the weight gain of *S. littoralis* caterpillars, and b) total seed production across 26 maternal half-sib families of *C. hirsuta* plants subjected to herbivory by *S. littoralis*. Grey lines represent maternal families’ averages in the constitutive (Control) state and in the induced state after JA addition the roots (JA). Black lines and dots represent overall averages across families.

### Effect of JA Treatment and Family Level Variations on GSL Production

The GSL profile of the *C. hirsuta* leaves consisted of 28 GSL compounds: 15 aliphatic-GSLs, 8 aromatic-GSLs, 3 indole-GSLs, and 2 unknown GSLs (Supplementary materials Table S1; Fig. 3a). We found that the maternal family background, but not the JA application, affected the multivariate GSL matrix in *C. hirsuta* leaves (Table 2, Fig. 3b). Specifically, maternal families explained 35% of the variance in the PERMANOVA, and such variation was also marginally explained by plant biomass (Table 2). We also found a maternal family effect for 16 out of the 28 GSLs (Table 1), a JA effect for five GSLs (GSL9: glucohirsutin, GSL12: 8-methylthiooctyl gsl, GSL14: hydroxymethylbutyl gsl, GSL17: veratryl gsl, GSL26: neoglucobrassicin; Table 1), and a biomass effect for four GSLs (GSL11: glucoberteroin, GSL24: glucobrassicin, GSL25: methoxyglucobrassicin, GSL26: neoglucobrassicin; Table 1). JA treatment significantly decreased the production of four out of those five compounds, except neoglucobrassicin, which increased its production by 25%. The production of GSL neoglucobrassicin was also significantly affected by plant biomass and the maternal families treatment, which explained 11% of the total variances (Table 1). We found no effect of JA treatment and maternal family on total levels of GSLs (Table 1). In addition, we found a significant interactive effect of family × JA (maternal family effect for induced production) for five GSLs (GSL1: glucoraphanin, GSL9: glucohirsutin, GSL10: glucoerucin, GSL13: gluconapoleiferin, GSL20: 5-benzoyloxypentyl) (Table S2).

**Table 2.** Permutational multivariate analysis of variance (PERMANOVA) table for testing the effect of JA treatment and family on the structure of the glucosinolate (GSLs) matrix.

| Factor | Df | MSQ | F value | R<sup>2</sup> | P value |
| --- | --- | --- | --- | --- | --- |
| JA treatment | 1 | 0.136213 | 2.04795 | 0.01494 | 0.118 |
| Family | 25 | 0.126121 | 1.89621 | 0.34592 | **0.005\*\*** |
| Plant biomass | 1 | 0.192319 | 2.89149 | 0.0211 | **0.054** |
| JA * Family | 25 | 0.063047 | 0.94791 | 0.17292 | 0.571 |
| Residuals | 61 | 0.066512 |  | 0.44512 |  |

Signif. codes: 0 '\*\*\*' 0.001 '\*\*' 0.01 '\*' 0.05 '.' 0.1 ' ' 1

**Fig. 3.**
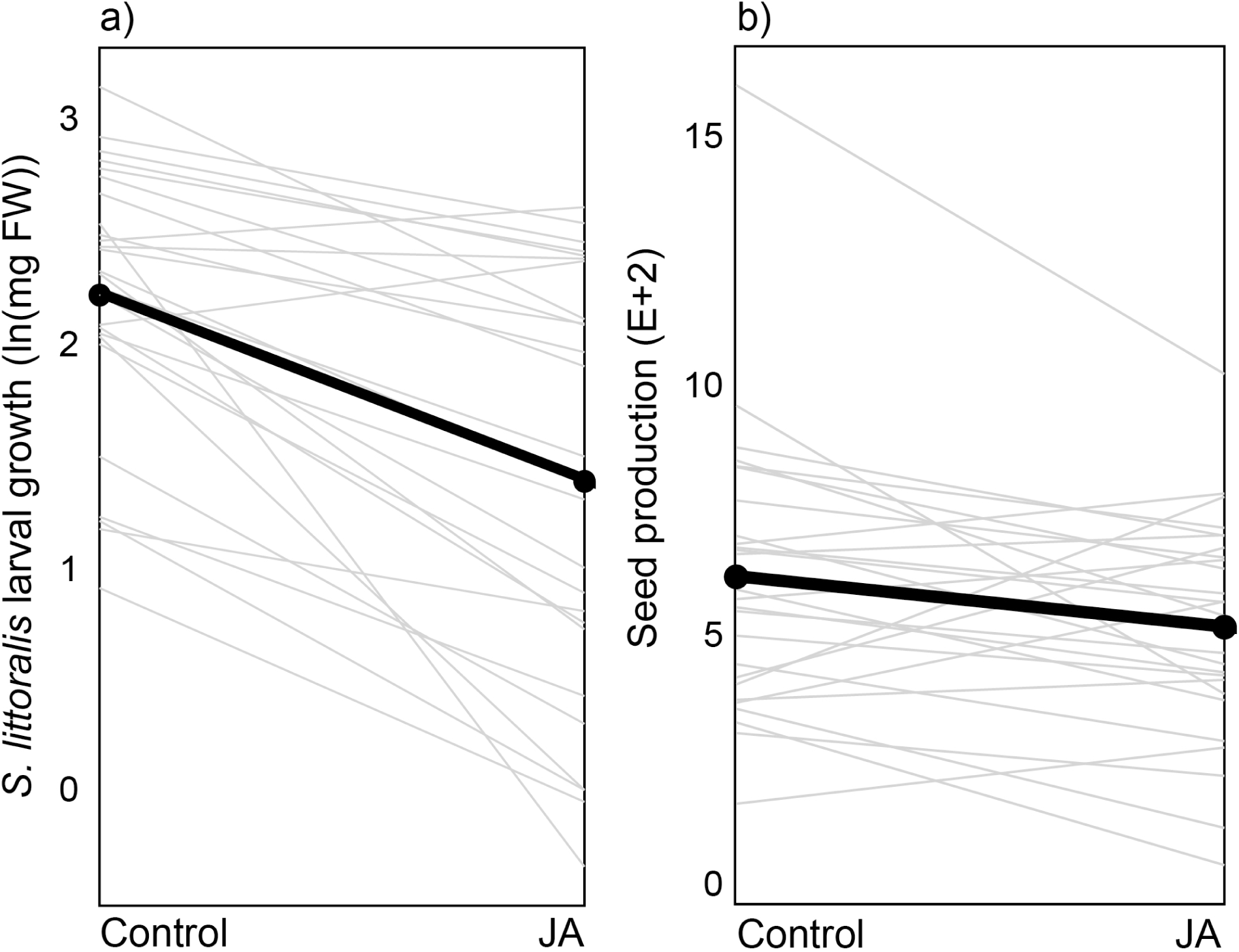
Glucosinolate (GSL) production across *Cardamine hirsuta* half-sib families. **a)** Barplot representation of the concentration of the individual GSLs in leaves of *C. hirsuta* plants that either received JA in the roots 4 days prior to the start of herbivory (JA, grey bars), or did not receive JA treatment in the roots (Control, open bars). Asterisks indicate a significant effect of JA treatment in production of GSLs. GSL1 = Glucoraphanin; GSL2 = Hydroxypropyl gsl; GSL3 = Progoitrin; GSL4 = Glucoalyssin; GSL5 = Glucoputranjivin; GSL6 = Gluconapin; GSL7 = Butyl gsl; GSL8 = Glucobrassicanapin; GSL9 = Glucohirsutin; GSL10 = Glucoerucin; GSL11 = Glucoberteroin; GSL12 = 8-Methylthiooctyl gsl; GSL13 = Gluconapoleiferin; GSL14 = Hydroxymethylbutyl gsl; GSL15 = 2-Methylbutyl gsl; GSL16 = Sinalbin; GSL17 = Veratryl gsl; GSL18 = Glucotropeolin; GSL19 = Trimethoxy gsl; GSL20 = 5-Benzoyloxypentyl; GSL21 = 2-Hydroxy-2-phenylethyl gsl; GSL22 = Gluconasturtiin; GSL23 = Hydroxybenzyl-methylether gsl; GSL24 = Glucobrassicin; GSL25 = Methoxyglucobrassicin; GSL26 = Neoglucobrassicin; GSL27 = Unknown.C16H23NO10S2; GSL28 = Unknown.C19H28N3O12S. **b)** Non-multidimensional scaling (nMDS) ordination of the individual glucosinolates found in *C. hirsuta* leaves across 26 plant families at the constitutive state (open dots), or after roots induction with JA (black dots). Numbers besides dots correspond to plant families.

### Effect of herbivory on Seed Set

Across all families, lifetime seed production in the control (no-herbivory) treatment was significantly higher compared to plants in induced and no-induction treatment that experienced herbivory (Fig. 4, *F*_*1,144*_ = 54.70, *p* <.0001). While *P. brassicae* and *S. littoralis* herbivory generally decreased seed set by 68% and 40%, respectively, we found strong genetic effect on seed set production after *S. littoralis* herbivory (Table 1, Fig. 2b). Finally, we found no significant JA treatment effect on seed set (Table 1, Fig. 4)

**Fig. 4.**
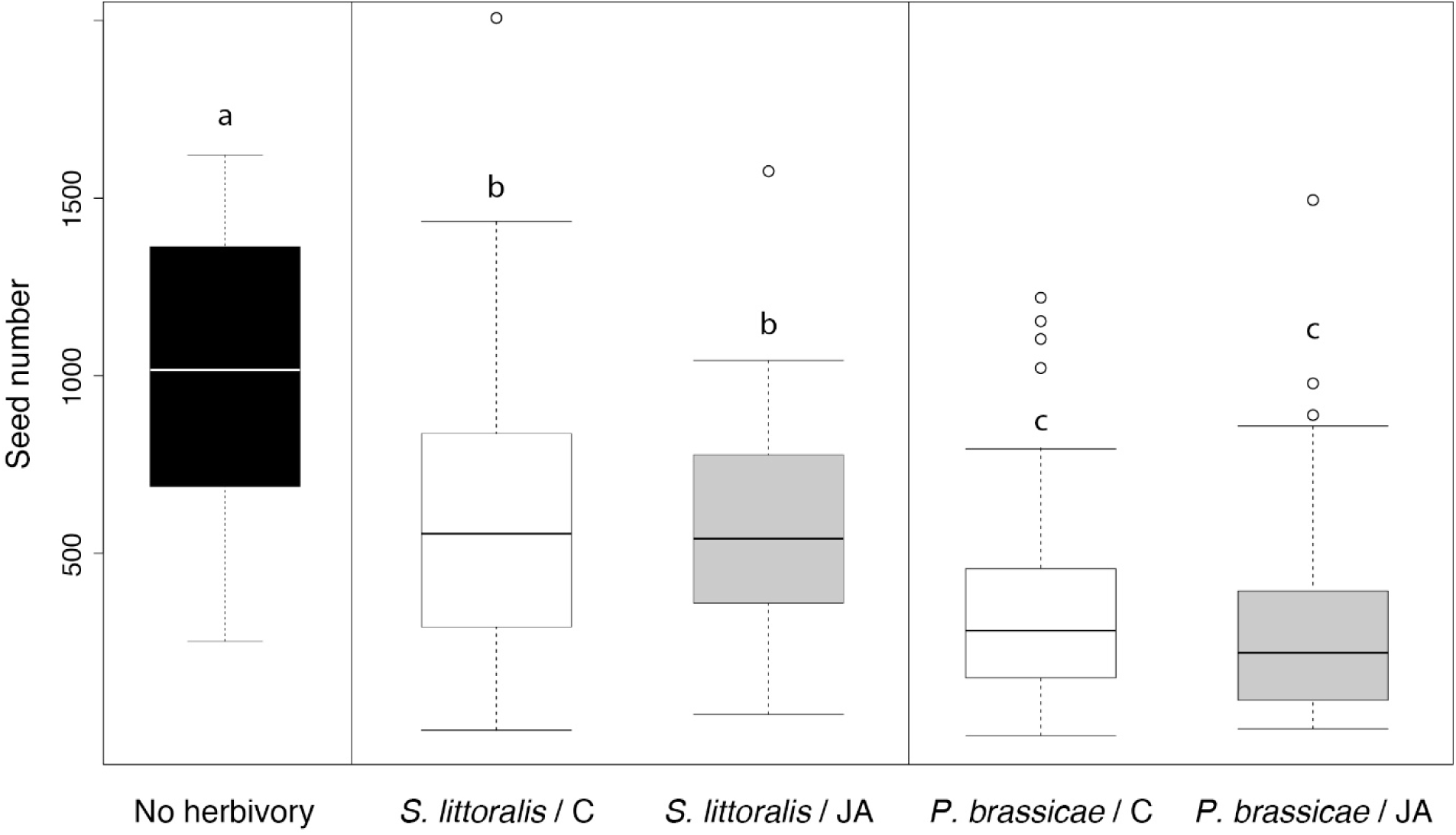
Herbivore impact on plant fitness. Boxplots show the total seed number produced by 26 maternal half-sib families of *C. hirsuta* plants in plants that did not receive JA in the roots nor treated aboveground herbivory (No herbivory), plants that received jasmonic acid (JA) in the roots 4 days prior to the start of herbivory by *Spodoptera littoralis* or *Pieris brassicae* (*P. brassicae* / JA, *S. littoralis* / JA), and plants subjected to herbivory either by *S. littoralis* or *P. brassicae* but that that were not treated by JA in the roots (*P. brassicae* / C, *S. littoralis* / C).

### Trade-offs Analyses

We detected a significant negative correlation (trade-off), between the constitutive production and the inducibility of total GSLs across all maternal families of *C. hirsuta* (*r* = - 0.82, *p* = 0.01, Fig. S1).

### Effect of JA Root Induction on Plant Fitness after Herbivore Attack

Mixed effect ANCOVA analyses showed that five GSLs (GSL4: glucoalyssin, GSL8: glucobrassicanapin, GSL10: glucoerucin, GSL11: glucoberteroin, GSL18: glucotropeolin), as well as the total GSLs production interacted with JA treatment for explaining seed production (Fig. 5, Table S3). In other words, JA induction changed the slope of the relationship between the GSLs and seed production from negative to neutral or even positive (Fig. 5). We also found marginally significant effect of JA×GSL for GSL16: sinalbin and GSL13: gluconapoleiferin.

**Fig. 5.**
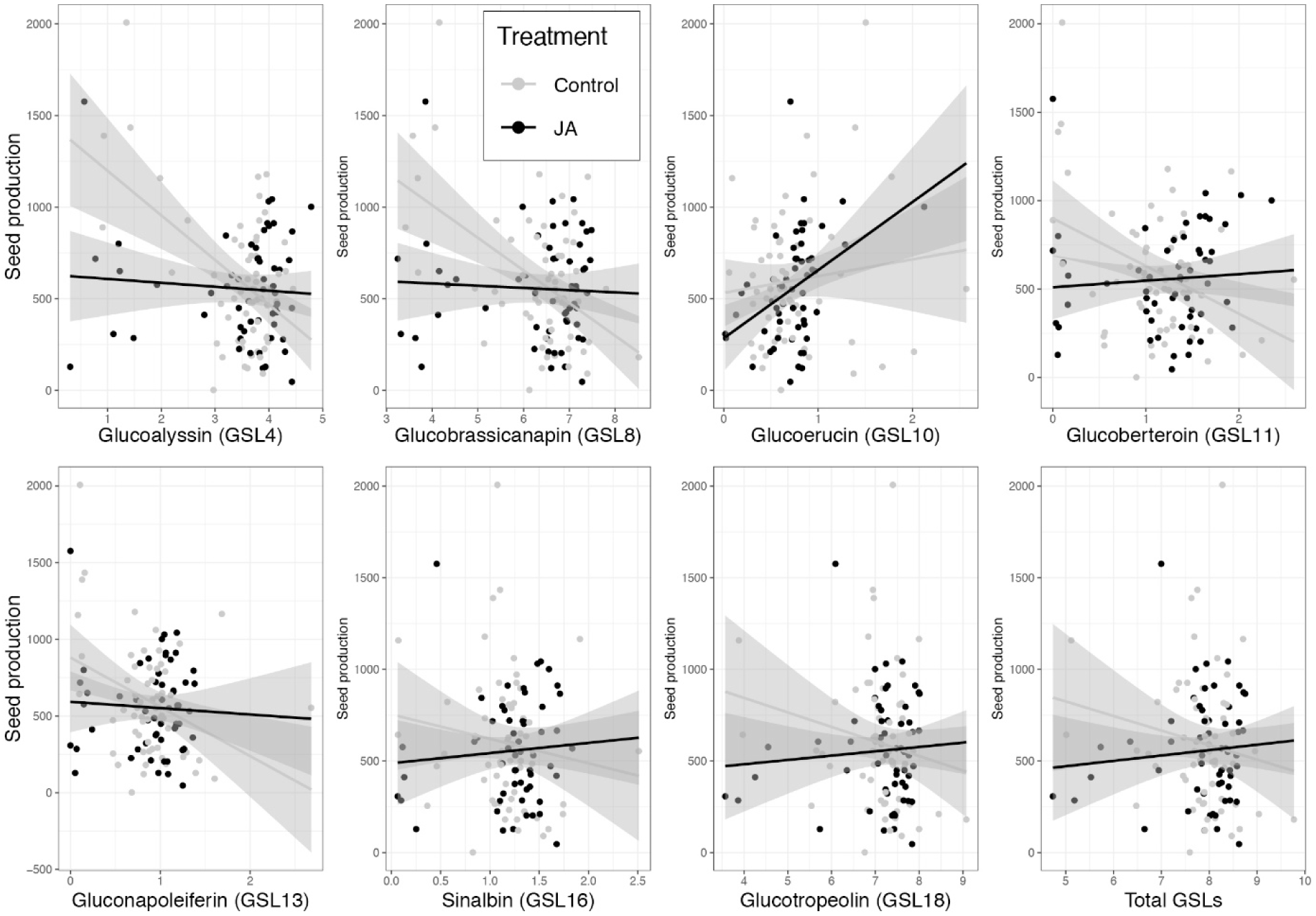
Fitness benefits of JA root induction during herbivory. Shown are interaction plots indicating the relationship between the total seed production and total glucosinolates, as well as seven individual GSLs that showed interaction with JA across all plants subjected to herbivory by *S. littoralis* in control and JA treatment (see Table S3).

## DISCUSSION

We found that the systemic induction, from below- to aboveground, of *C. hirsuta* plants significantly decreased the weight gain of a generalist leaf chewing herbivore, but such effect was highly variable across plant maternal families. Analyses of plants’ secondary metabolites showed that JA root induction affected the production of several GSLs aboveground, and significantly ameliorated plant fitness after leaf chewing herbivore attack. Below, we discuss the implications of these findings for the ecology and evolution of plant defense against herbivores in natural brassicaceaous systems.

### Effect of Root JA Treatment on Insect Resistance and Aboveground Glucosinolate Production

One of the principal results of our study is that the JA root application increased resistance against the generalist herbivore (*S. littoralis*), while JA application had no effect on the specialist herbivore (*P. brassicae*). These results are in line with several previous studies (Bodenhausen and Reymond 2007; Giamoustaris and Mithen 1995; Lankau 2007; Raybould and Moyes 2001). For instance, root JA application to *Brassica oleracea* roots resulted in reduced weight gain of generalist herbivore *Mamestra brassicae*, whereas the specialist *P. rapae* was unaffected (van Dam and Oomen 2008), and root induction resulted in more infestation by AG specialists in field-grown *B. oleracea* plants (Pierre et al. 2013). Indeed, specialist herbivores of the Brassicaceae not only tolerate GSLs, they also utilize these compounds in host recognition (Raybould and Moyes 2001). On the other hand, the negative effect of GSLs on generalist herbivore performance has also been confirmed in previous studies (Rasmann et al. 2015; Schlaeppi et al. 2008; Schweiger et al. 2014; Schweizer et al. 2017), which confirms strong context-dependency in plant-herbivore interaction.

Contrary to general expectations, we did not detect differences in the production of total GSLs between control and JA-treated plants, and found that the GSL production in leaves was related by a large extend to plant biomass, a common phenomenon when studying secondary metabolite production in plants (Glynn et al. 2003; Traw 2002; Züst et al. 2015). Although previous works with Brassicaceae plants showed that BG herbivory, or root induction by JA, increases total levels of GSL in shoots (Griffiths et al. 1994; Pierre et al. 2012; van Dam and Oomen 2008; van Dam et al. 2004), similarly to this one, other studies failed to detect such changes in production of total GSLs (Agrawal 2000; Pierre et al. 2013). These results indicate that the systemic induced responses in plants from BG to AG can be species (as well as genotypic, such as in this study) specific, and that uniquely measuring the total amount can often be misleading in plant-herbivore interaction studies..

Changes in phytochemical diversity in response to induction is, on the other hand, likely a more important component of plant defense against herbivory (Agrawal 2000; Berenbaum and Zangerl 1996; Lindig-Cisneros et al. 1997). Accordingly, the results of our multivariate analysis showed that among the five families that are distinctive with respect to their GSL profiles (Fig. 1b.), two families exhibited greater resistance against *S. littoralis* (family 5 & 7). In fact, family 7, which showed the most distinctive GSL composition in the NMDS, was the most responsive family to JA treatment in terms of inducibility of overall GSLs and the most-resistant family against herbivory by *S. littoralis*. In addition, in contrast to total GSL levels, we observed that an indolic GSL, neoglucobrassicin, is the only compound that was both significantly induced by JA (Table 1) and also negatively correlated with *S. littoralis* weight gain (results not shown, but results from linear mixed model for testing the interactive effect of the JA treatment and neoglucobrassicin production on *S. littoralis* weight gain: JA effect; F_x,y_ = 18.34, p < 0.001; neoglucobrassicin effect; F_x,y_ = 5.34, p = 0.02; and their interaction: F_x,y_ = 3.46, P = 0.07). Indole GSLs have been shown to be induced by herbivory and to affect the growth and development of insect herbivores in other systems (Irwin et al. 2003; Rostás et al. 2002). Selective induction of indolic GSLs have been reported in *B. napus, B. rapae* and *B. juncea* in response to herbivory by flea beetles (Bodnaryk 1992). For instance, the concentration of neoglucobrassicin was increased considerably in leaves of *B. napus* as a result of topical application of methyl JA to aerial part of the plant (Doughty et al. 1995), as well as in *B. rapae* and *B. napus* plants treated with specialist herbivores (Koritsas et al. 1991; Rostás et al. 2002). The same pattern of induction of neoglucobrassicin was observed in the roots of *B. napus* that were damaged by *Delia floralis* root maggots (Hopkins et al. 1998). In another study, the only compound that was shown to affect the performance of *P. rapae* feeding on *B. oleracea* plants was neoglucobrassicin (Harvey et al. 2007).. Together, these results suggest that the total amount of GSLs in Brassicaceaous plants can often be misleading when predicting plant resistance, while, on the other hand, individual GSLs bear differential toxicities might be better predictors of plant resistance.

### Does Below-to-Aboveground Systemic Induction of GSLs Affect Plant Fitness?

Demonstrating the effect of induced response on plant fitness is crucial for documenting that they truly serve as a defensive response (Erb 2018). We found that herbivory, overall, decreased plant fitness (seed production) by more than 50%, clearly confirming the well-documented negative consequence of herbivory on plant fitness (Agrawal 1998; Agrawal 1999; Kessler and Baldwin 2004; Maron 1998; Mothershead and Marquis 2000). If herbivory decreases plant fitness and plants possess genetic variation for traits affecting herbivory and enhancing fitness, then herbivores may act as strong selective agents for more resistant plants by promoting inducibility of specific toxic molecules. Accordingly, in *C. hirsuta*, we showed that root JA-mediated induced systemic production of seven GSL compounds in shoots increases seed production in plants exposed to shoot herbivory, compared to plants that did not received JA treatment. This fitness impact has important implications. First, inducible systemic resistance may be an example of adaptive plasticity in plants. Adaptive plasticity is defined by the higher fitness of individuals expressing different phenotypes in a particular environment (Vijendravarma et al. 2015). Thus, the induction of GSLs compounds after root damage can be seen as an adaptive plastic response for *C. hirsuta* plants (Agrawal 1999; Agrawal 2000). Nonetheless, to be fully convincing, arguments about adaptive plastic responses should also be linked to ecological setting. In this case, we could speculate that *C. hirsuta* plants are likely damaged in their roots, by e.g. root fly maggots, every spring before pierids or other generalist butterflies start feeding on these plants. To date, due to obvious methodological limitations of measuring rates of root herbivory in the field, we only have anecdotal information on the timing and amount of root damage in natural systems (Johnson and Rasmann 2015). For now we can only speculate that the observed genetic variation in inducibility from below to aboveground is shaped by the natural variation in root and shoot herbivory.

The second implication of our fitness-related results concerns the evolution of the systemic response from root to shoots. In order for such a trait to evolve by natural selection, there must be heritable variation that affects fitness. We detected genetic variation in induced production of five GSL compounds (significant interactive family ×JA effect). Within these five GSLs, two compounds (GSL10 and 13) were among the seven individual compounds found to be positively affecting seed set when induced by JA. In other words, plant families possessing the ability for increased production of these seven compounds in the induced state could hinder the negative fitness effect of herbivory.

Finally, genetic variation in inducibility could also have been maintained by physiological trade-offs. Accordingly, we showed that the inducibility of total GSLs and neoglucobrassicin negatively correlated with constitutive investment in both traits. It is generally assumed that constitutive and induced defenses should trade off, as the anti-herbivore defenses are costly for plants (Karban and Baldwin 1997; Karban and Myers 1989; Zangerl and Bazzaz 1992). Thus, most *C. hirsuta* families employ economy in direct chemical defense production, by favoring either a constitutive or an inducible strategy. Altogether, ecological and physiological trade-offs may contribute in maintaining the necessary genetic variation in inducibility of specific GSLs, ultimately generating the raw material for selection to act upon.

## Author contributions

MB and SR conceived and designed the experiments. MB conducted experiments and chemical analyses. MB and SR analysed the data and wrote the manuscript. The authors declare no conflicts of interest.

## Acknowledgements

We thank Mégane Rohrer and Ludovico Formenti for assisting with experimental work and trait measurements. This work was supported by Swiss National Science Foundation grants 179481 and 159869 to SR.

