## Supplementary Tables and Figures for "Genotypic Variation in Below-to Aboveground Systemic Induction of Glucosinolates Mediates Plant Fitness Consequences under Herbivore Attack"

*Corresponding author

***Authors’ information****:*

**Table S1.** Family mean raw values for concentrations of individual glucosinolates in plants treated with JA in the roots and control plants (no-induction) across 26 *C. hirsuta* plant families. (Separate .excel file). GSL1 = Glucoraphanin; GSL2 = Hydroxypropyl gsl; GSL3 = Progoitrin; GSL4 = Glucoalyssin; GSL5 = Glucoputranjivin; GSL6 = Gluconapin; GSL7 = Butyl gsl; GSL8 = Glucobrassicanapin; GSL9 = Glucohirsutin; GSL10 = Glucoerucin; GSL11 = Glucoberteroin; GSL12 = 8-Methylthiooctyl gsl ; GSL13 = Gluconapoleiferin; GSL14 = Hydroxymethylbutyl gsl ; GSL15 = 2-Methylbutyl gsl; GSL16 = Sinalbin; GSL17 = Veratryl gsl; GSL18 = Glucotropeolin; GSL19 = Trimethoxy gsl; GSL20 = 5-Benzoyloxypentyl ; GSL21 = 2-Hydroxy-2-phenylethyl gsl; GSL22 = Gluconasturtiin; GSL23 = Hydroxybenzyl-methylether gsl; GSL24 = Glucobrassicin; GSL25 = Methoxyglucobrassicin; GSL26 = Neoglucobrassicin; GSL27 = Unknown.C16H23NO10S2; GSL28 = Unknown.C19H28N3O12S3.

**Table S2.** Mixed effect model table for testing the effect of JA induction treatment in the roots of *Cardamine hirsuta* plants, maternal families, and their biomass on individual and total glucosinolates (GSL*), as well as the interaction between JA treatment and maternal families. *C. hirsuta* plant families nested within populations was the fixed factor. Significant interactive effects are marked in bold and indicate a significant family effect of induction.

| **GSL†** | **Factor** | **SSq** | **MSq** | **F value** | **Pr(>F)** |  |
| --- | --- | --- | --- | --- | --- | --- |
| **GSL1** | JA | 0.434 | 0.434 | 1.984 | 0.164 |  |
|  | Fam | 10.192 | 0.4077 | 1.863 | 0.023 | * |
|  | Plant.biomass | 0.538 | 0.5382 | 2.46 | 0.122 |  |
|  | **JA:Fam** | **9.347** | **0.3739** | **1.709** | **0.044** | ***** |
| GSL2 | JA | 0.115 | 0.11548 | 1.242 | 0.269 |  |
|  | Fam | 1.919 | 0.07675 | 0.826 | 0.697 |  |
|  | Plant biomass | 0.061 | 0.06071 | 0.653 | 0.422 |  |
|  | JA:Fam | 1.347 | 0.0539 | 0.58 | 0.934 |  |
| GSL3 | JA | 0.105 | 0.1045 | 0.746 | 0.391 |  |
|  | Fam | 10.265 | 0.4106 | 2.932 | 0.0002 | *** |
|  | Plant biomass | 0.403 | 0.4034 | 2.881 | 0.095 | . |
|  | JA:Fam | 4.012 | 0.1605 | 1.146 | 0.323 |  |
| GSL4 | JA | 0 | 0.0022 | 0.007 | 0.936 |  |
|  | Fam | 67.34 | 2.6936 | 7.813 | 0 | *** |
|  | Plant biomass | 0.17 | 0.167 | 0.484 | 0.489 |  |
|  | JA:Fam | 7.97 | 0.3186 | 0.924 | 0.573 |  |
| GSL5 | JA | 0.02 | 0.0202 | 0.07 | 0.792 |  |
|  | Fam | 13.295 | 0.5318 | 1.856 | 0.024 | * |
|  | Plant biomass | 0.008 | 0.0077 | 0.027 | 0.87 |  |
|  | JA:Fam | 3.553 | 0.1421 | 0.496 | 0.973 |  |
| GSL6 | JA | 0.02 | 0.0157 | 0.017 | 0.896 |  |
|  | Fam | 19.74 | 0.7897 | 0.861 | 0.652 |  |
|  | Plant biomass | 0.01 | 0.0068 | 0.007 | 0.931 |  |
|  | JA:Fam | 14.85 | 0.594 | 0.648 | 0.886 |  |
| GSL7 | JA | 0.007 | 0.0066 | 0.023 | 0.881 |  |
|  | Fam | 13.696 | 0.5478 | 1.903 | 0.02 | * |
|  | Plant biomass | 0.006 | 0.006 | 0.021 | 0.886 |  |
|  | JA:Fam | 3.849 | 0.154 | 0.535 | 0.958 |  |
| GSL8 | JA | 0 | 0.0003 | 0 | 0.986 |  |
|  | Fam | 76.52 | 3.0607 | 3.573 | 0 | *** |
|  | Plant biomass | 0.02 | 0.0185 | 0.022 | 0.884 |  |
|  | JA:Fam | 15.95 | 0.638 | 0.745 | 0.791 |  |
| **GSL9** | JA | 0.2331 | 0.23312 | 200.61 | 0 | *** |
|  | Fam | 2.9661 | 0.11864 | 102.1 | 0 | *** |
|  | Plant biomass | 0.0154 | 0.01538 | 13.23 | 0.0005 | *** |
|  | **JA:Fam** | **2.2951** | **0.0918** | **79** | **0** | ******* |
| **GSL10** | JA | 0.012 | 0.01185 | 0.108 | 0.744 |  |
|  | Fam | 5.944 | 0.23775 | 2.164 | 0.007 | ** |
|  | Plant biomass | 0.314 | 0.31408 | 2.859 | 0.095 | . |
|  | **JA:Fam** | **5.028** | **0.20112** | **1.831** | **0.026** | ***** |
| GSL11 | JA | 0.187 | 0.1871 | 1.075 | 0.304 |  |
|  | Fam | 18.3 | 0.732 | 4.206 | 0 | *** |
|  | Plant biomass | 0.355 | 0.3553 | 2.041 | 0.158 |  |
|  | JA:Fam | 5.025 | 0.201 | 1.155 | 0.314 |  |
| GSL12 | JA | 0.273 | 0.27267 | 3.557 | 0.064 | . |
|  | Fam | 3.178 | 0.12713 | 1.658 | 0.053 | . |
|  | Plant biomass | 0.075 | 0.07451 | 0.972 | 0.328 |  |
|  | JA:Fam | 2.754 | 0.11016 | 1.437 | 0.122 |  |
| **GSL13** | JA | 0.026 | 0.0255 | 0.301 | 0.585 |  |
|  | Fam | 9.557 | 0.3823 | 4.505 | 0 | *** |
|  | Plant biomass | 0.082 | 0.0822 | 0.969 | 0.329 |  |
|  | **JA:Fam** | **3.479** | **0.1391** | **1.64** | **0.057** | **.** |
| GSL14 | JA | 0.382 | 0.382 | 5.251 | 0.025 | * |
|  | Fam | 2.915 | 0.1166 | 1.603 | 0.066 | . |
|  | Plant biomass | 0.013 | 0.0135 | 0.185 | 0.669 |  |
|  | JA:Fam | 2.474 | 0.099 | 1.36 | 0.161 |  |
| GSL15 | JA | 0.198 | 0.19795 | 1.072 | 0.304 |  |
|  | Fam | 4.938 | 0.19753 | 1.07 | 0.4 |  |
|  | Plant biomass | 0.009 | 0.00893 | 0.048 | 0.827 |  |
|  | JA:Fam | 4.933 | 0.19732 | 1.069 | 0.401 |  |
| GSL16 | JA | 0.126 | 0.12622 | 0.929 | 0.34 |  |
|  | Fam | 5.281 | 0.21126 | 1.555 | 0.079 | . |
|  | Plant biomass | 0.013 | 0.01257 | 0.093 | 0.762 |  |
|  | JA:Fam | 4.345 | 0.17381 | 1.279 | 0.212 |  |
| GSL17 | JA | 0.268 | 0.26768 | 3.45 | 0.068 | . |
|  | Fam | 3.129 | 0.12518 | 1.613 | 0.063 | . |
|  | Plant biomass | 0.072 | 0.07241 | 0.933 | 0.338 |  |
|  | JA:Fam | 2.737 | 0.10947 | 1.411 | 0.135 |  |
| GSL18 | JA | 0.01 | 0.0131 | 0.014 | 0.905 |  |
|  | Fam | 18.47 | 0.7388 | 0.81 | 0.716 |  |
|  | Plant biomass | 0 | 0.0049 | 0.005 | 0.942 |  |
|  | JA:Fam | 18.35 | 0.734 | 0.805 | 0.722 |  |
| GSL19 | JA | 0.26 | 0.25955 | 3.297 | 0.074 | . |
|  | Fam | 3.123 | 0.12493 | 1.587 | 0.07 | . |
|  | Plant biomass | 0.074 | 0.07416 | 0.942 | 0.335 |  |
|  | JA:Fam | 2.733 | 0.10934 | 1.389 | 0.146 |  |
| **GSL20** | JA | 0.0767 | 0.07665 | 3.255 | 0.08 | . |
|  | Fam | 2.0809 | 0.08324 | 3.535 | 0 | *** |
|  | Plant biomass | 0.044 | 0.04401 | 1.869 | 0.176 |  |
|  | **JA:Fam** | **2.0598** | **0.08239** | **3.499** | **0** | ******* |
| GSL21 | JA | 0.273 | 0.27292 | 3.686 | 0.06 | . |
|  | Fam | 3.22 | 0.1288 | 1.74 | 0.04 | * |
|  | Plant biomass | 0.009 | 0.00906 | 0.122 | 0.728 |  |
|  | JA:Fam | 2.072 | 0.08288 | 1.119 | 0.348 |  |
| GSL22 | JA | 0.1 | 0.1048 | 0.187 | 0.667 |  |
|  | Fam | 49.09 | 1.9634 | 3.51 | 0 | *** |
|  | Plant biomass | 0.01 | 0.0141 | 0.025 | 0.874 |  |
|  | JA:Fam | 14.01 | 0.5605 | 1.002 | 0.478 |  |
| GSL23 | JA | 0.273 | 0.27292 | 3.686 | 0.05 | . |
|  | Fam | 3.22 | 0.1288 | 1.74 | 0.04 | * |
|  | Plant biomass | 0.009 | 0.00906 | 0.122 | 0.728 |  |
|  | JA:Fam | 2.072 | 0.08288 | 1.119 | 0.348 |  |
| GSL24 | JA | 0.167 | 0.1667 | 0.436 | 0.511 |  |
|  | Fam | 17.74 | 0.7096 | 1.857 | 0.024 | * |
|  | Plant biomass | 0.991 | 0.9906 | 2.592 | 0.112 |  |
|  | JA:Fam | 7.189 | 0.2875 | 0.752 | 0.783 |  |
| GSL25 | JA | 0.01 | 0.0105 | 0.029 | 0.866 |  |
|  | Fam | 32.19 | 1.2878 | 3.491 | 0 | *** |
|  | Plant biomass | 2.06 | 2.0609 | 5.587 | 0.021 | * |
|  | JA:Fam | 5.06 | 0.2023 | 0.548 | 0.952 |  |
| GSL26 | JA | 0.0429 | 0.04295 | 3.247 | 0.07 | . |
|  | Fam | 0.8781 | 0.03512 | 2.655 | 0.0008 | *** |
|  | Plant biomass | 0.0411 | 0.0411 | 3.107 | 0.08 | . |
|  | JA:Fam | 0.3212 | 0.01285 | 0.971 | 0.514 |  |
| GSL27 | JA | 0.245 | 0.24525 | 2.996 | 0.088 | . |
|  | Fam | 3.133 | 0.12534 | 1.531 | 0.087 | . |
|  | Plant biomass | 0.027 | 0.02651 | 0.324 | 0.571 |  |
|  | JA:Fam | 2.242 | 0.08967 | 1.095 | 0.373 |  |
| GSL28 | JA | 0.379 | 0.3792 | 4.906 | 0.03 | * |
|  | Fam | 3.122 | 0.1249 | 1.615 | 0.063 | . |
|  | Plant biomass | 0.015 | 0.0153 | 0.198 | 0.658 |  |
|  | JA:Fam | 2.172 | 0.0869 | 1.124 | 0.344 |  |
| GSL Total | JA | 0.03 | 0.0252 | 0.034 | 0.855 |  |
|  | Fam | 15.22 | 0.6089 | 0.816 | 0.708 |  |
|  | Plant biomass | 0 | 0.0009 | 0.001 | 0.973 |  |
|  | JA:Fam | 14.52 | 0.5806 | 0.778 | 0.753 |  |

Signif. codes: *** < 0.001,** <0.01, * <0.05, ° <0.1

**†** GSL1 = Glucoraphanin; GSL2 = Hydroxypropyl gsl; GSL3 = Progoitrin; GSL4 = Glucoalyssin; GSL5 = Glucoputranjivin; GSL6 = Gluconapin; GSL7 = Butyl gsl; GSL8 = Glucobrassicanapin; GSL9 = Glucohirsutin; GSL10 = Glucoerucin; GSL11 = Glucoberteroin; GSL12 = 8-Methylthiooctyl gsl ; GSL13 = Gluconapoleiferin; GSL14 = Hydroxymethylbutyl gsl ; GSL15 = 2-Methylbutyl gsl; GSL16 = Sinalbin; GSL17 = Veratryl gsl; GSL18 = Glucotropeolin; GSL19 = Trimethoxy gsl; GSL20 = 5-Benzoyloxypentyl ; GSL21 = 2-Hydroxy-2-phenylethyl gsl; GSL22 = Gluconasturtiin; GSL23 = Hydroxybenzyl-methylether gsl; GSL24 = Glucobrassicin; GSL25 = Methoxyglucobrassicin; GSL26 = Neoglucobrassicin; GSL27 = Unknown.C16H23NO10S2; GSL28 = Unknown.C19H28N3O12S3.

**Table S3.** Mixed effect model table for testing the effect of individual and total glucosinolates (GSL*), and JA induction treatment in the roots on *Cardamine hirsuta* plants **lifetime seed production**. *C. hirsuta* plant families nested within populations was the random factor. Significant interactive effects are marked in bold and indicate a significant positive effect of induction on plant fitness.

| **GSL** | **Factor** | **SSQ** | **DenDF** | **F** | **Pr(>F)** |  |
| --- | --- | --- | --- | --- | --- | --- |
| GSL1 | GSL | 1156228 | 110.42 | 15.577 | 0 | *** |
|  | JA | 7932 | 101.06 | 0.107 | 0.744 |  |
|  | GSL:JA | 41180 | 107.36 | 0.555 | 0.458 |  |
| GSL2 | GSL | 106983 | 108.405 | 1.401 | 0.239 |  |
|  | JA | 151646 | 92.382 | 1.986 | 0.162 |  |
|  | GSL:JA | 97516 | 108.566 | 1.277 | 0.261 |  |
| GSL3 | GSL | 58413 | 110.45 | 0.751 | 0.388 |  |
|  | JA | 71022 | 97.437 | 0.914 | 0.342 |  |
|  | GSL:JA | 2960 | 100.234 | 0.038 | 0.846 |  |
| **GSL4** | GSL | 33932 | 111.383 | 0.468 | 0.495 |  |
|  | JA | 606202 | 93.809 | 8.363 | 0.005 | ** |
|  | **GSL:JA** | **472550** | **95.701** | **6.519** | **0.012** | ***** |
| GSL5 | GSL | 10066 | 106.243 | 0.133 | 0.716 |  |
|  | JA | 226815 | 94.149 | 2.995 | 0.087 | . |
|  | GSL:JA | 104181 | 94.983 | 1.376 | 0.244 |  |
| GSL6 | GSL | 118812 | 101.762 | 1.563 | 0.214 |  |
|  | JA | 172382 | 99.083 | 2.268 | 0.135 |  |
|  | GSL:JA | 68980 | 100.626 | 0.907 | 0.343 |  |
| GSL7 | GSL | 11793 | 109.586 | 0.156 | 0.694 |  |
|  | JA | 241666 | 94.647 | 3.187 | 0.077 | . |
|  | GSL:JA | 113842 | 94.788 | 1.501 | 0.224 |  |
| **GSL8** | GSL | 1358 | 106.779 | 0.019 | 0.891 |  |
|  | JA | 620941 | 98.762 | 8.699 | 0.004 | ** |
|  | **GSL:JA** | **508015** | **100.846** | **7.117** | **0.009** | ****** |
| GSL9 | GSL | 71423 | 96.483 | 0.92 | 0.34 |  |
|  | JA | 211755 | 93.888 | 2.727 | 0.102 |  |
|  | GSL:JA | 74183 | 96.393 | 0.955 | 0.331 |  |
| **GSL10** | GSL | 448846 | 103.661 | 6.181 | 0.015 | * |
|  | JA | 437582 | 97.152 | 6.026 | 0.016 | * |
|  | **GSL:JA** | **357717** | **105.137** | **4.926** | **0.029** | ***** |
| **GSL11** | GSL | 110965 | 109.472 | 1.642 | 0.203 |  |
|  | JA | 866668 | 95.148 | 12.828 | 0.001 | *** |
|  | **GSL:JA** | **731786** | **98.86** | **10.831** | **0.001** | ****** |
| GSL12 | GSL | 93884 | 108.2 | 1.229 | 0.27 |  |
|  | JA | 161440 | 92.51 | 2.113 | 0.15 |  |
|  | GSL:JA | 105176 | 108.43 | 1.376 | 0.243 |  |
| **GSL13** | GSL | 41844 | 109.56 | 0.564 | 0.454 |  |
|  | JA | 335753 | 101.06 | 4.521 | 0.036 | * |
|  | **GSL:JA** | **208650** | **103.9** | **2.81** | **0.097** | **.** |
| GSL14 | GSL | 1392 | 112 | 0.018 | 0.894 |  |
|  | JA | 89371 | 98.968 | 1.152 | 0.286 |  |
|  | GSL:JA | 1261 | 111.999 | 0.016 | 0.899 |  |
| GSL15 | GSL | 7293 | 106.896 | 0.094 | 0.76 |  |
|  | JA | 168913 | 94.224 | 2.182 | 0.143 |  |
|  | GSL:JA | 33672 | 107.872 | 0.435 | 0.511 |  |
| **GSL16** | GSL | 57931 | 103.091 | 0.781 | 0.379 |  |
|  | JA | 348929 | 99.687 | 4.703 | 0.033 | * |
|  | **GSL:JA** | **226158** | **101.247** | **3.048** | **0.084** | **.** |
| GSL17 | GSL | 102615 | 108.291 | 1.345 | 0.249 |  |
|  | JA | 165506 | 92.558 | 2.169 | 0.144 |  |
|  | GSL:JA | 114380 | 108.541 | 1.499 | 0.224 |  |
| **GSL18** | GSL | 15732 | 103.067 | 0.213 | 0.646 |  |
|  | JA | 425820 | 99.474 | 5.759 | 0.018 | * |
|  | **GSL:JA** | **300046** | **100.707** | **4.058** | **0.047** | ***** |
| GSL19 | GSL | 91743 | 108.328 | 1.199 | 0.276 |  |
|  | JA | 159375 | 92.572 | 2.083 | 0.152 |  |
|  | GSL:JA | 102318 | 108.591 | 1.337 | 0.25 |  |
| GSL20 | GSL | 30342 | 107.748 | 0.392 | 0.533 |  |
|  | JA | 177466 | 95.738 | 2.294 | 0.133 |  |
|  | GSL:JA | 64822 | 109.915 | 0.838 | 0.362 |  |
| GSL21 | GSL | 48213 | 107.02 | 0.628 | 0.43 |  |
|  | JA | 172833 | 103.4 | 2.25 | 0.137 |  |
|  | GSL:JA | 49765 | 106.75 | 0.648 | 0.423 |  |
| GSL22 | GSL | 143028 | 55.071 | 1.848 | 0.18 |  |
|  | JA | 131686 | 94.842 | 1.702 | 0.195 |  |
|  | GSL:JA | 11486 | 92.213 | 0.148 | 0.701 |  |
| GSL23 | GSL | 48213 | 107.02 | 0.628 | 0.43 |  |
|  | JA | 172833 | 103.4 | 2.25 | 0.137 |  |
|  | GSL:JA | 49765 | 106.75 | 0.648 | 0.423 |  |
| GSL24 | GSL | 75273 | 107.347 | 0.99 | 0.322 |  |
|  | JA | 184255 | 96.422 | 2.423 | 0.123 |  |
|  | GSL:JA | 72823 | 98.054 | 0.958 | 0.33 |  |
| GSL25 | GSL | 1295 | 110.232 | 0.017 | 0.897 |  |
|  | JA | 28640 | 93.685 | 0.37 | 0.544 |  |
|  | GSL:JA | 12286 | 94.145 | 0.159 | 0.691 |  |
| GSL26 | GSL | 53148 | 111.984 | 0.686 | 0.409 |  |
|  | JA | 68736 | 94.376 | 0.887 | 0.349 |  |
|  | GSL:JA | 2783 | 92.884 | 0.036 | 0.85 |  |
| GSL27 | GSL | 78493 | 108.242 | 1.036 | 0.311 |  |
|  | JA | 182841 | 92.393 | 2.413 | 0.124 |  |
|  | GSL:JA | 125252 | 108.49 | 1.653 | 0.201 |  |
| GSL28 | GSL | 24322 | 101.51 | 0.313 | 0.577 |  |
|  | JA | 37445 | 97.862 | 0.481 | 0.49 |  |
|  | GSL:JA | 24054 | 101.532 | 0.309 | 0.579 |  |
| **GSL Total** | GSL | 26563 | 102.043 | 0.362 | 0.549 |  |
|  | JA | 461821 | 99.455 | 6.294 | 0.014 | * |
|  | **GSL:JA** | **332386** | **100.822** | **4.53** | **0.036** | ***** |

Signif. codes: *** < 0.001,** <0.01, * <0.05, ° <0.1

* GSL1 = Glucoraphanin; GSL2 = Hydroxypropyl gsl; GSL3 = Progoitrin; GSL4 = Glucoalyssin; GSL5 = Glucoputranjivin; GSL6 = Gluconapin; GSL7 = Butyl gsl; GSL8 = Glucobrassicanapin; GSL9 = Glucohirsutin; GSL10 = Glucoerucin; GSL11 = Glucoberteroin; GSL12 = 8-Methylthiooctyl gsl ; GSL13 = Gluconapoleiferin; GSL14 = Hydroxymethylbutyl gsl ; GSL15 = 2-Methylbutyl gsl; GSL16 = Sinalbin; GSL17 = Veratryl gsl; GSL18 = Glucotropeolin; GSL19 = Trimethoxy gsl; GSL20 = 5-Benzoyloxypentyl ; GSL21 = 2-Hydroxy-2-phenylethyl gsl; GSL22 = Gluconasturtiin; GSL23 = Hydroxybenzyl-methylether gsl; GSL24 = Glucobrassicin; GSL25 = Methoxyglucobrassicin; GSL26 = Neoglucobrassicin; GSL27 = Unknown.C16H23NO10S2; GSL28 = Unknown.C19H28N3O12S3.

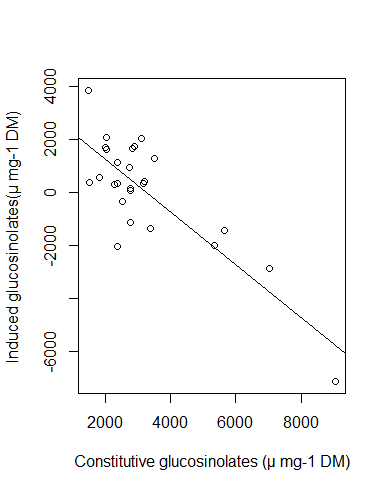

**Figure S1.** Trade-off between constitutive and inducibility in production of total glucosinolates in the shoots of *C. hirsuta* plants. Inducibility is measured as the difference between JA-treated and control values for each traits.
